# Metatranscriptomics reveals a shift in microbial community composition and function during summer months in a coastal marine environment

**DOI:** 10.1101/2022.01.21.477286

**Authors:** Ben J. G. Sutherland, Jan F. Finke, Robert Saunders, Snehal Warne, Angela D. Schulze, Jeff H. T. Strohm, Amy M. Chan, Curtis A. Suttle, Kristina M. Miller

## Abstract

Temperate coastal marine waters are often thermally stratified from spring through fall but can be dynamic and disrupted by tidal currents and wind-driven upwelling. These mixing events introduce deeper, cooler water with a higher partial pressure of CO2 (pCO_2_), and its associated microbial communities to the surface. Anecdotally, these events impact shellfish hatcheries and farms, warranting improved understanding of changes in composition and activity of marine microbial communities in relation to environmental processes. To characterize both compositional and functional changes associated with abiotic factors, here we generate a reference metatranscriptome from the Strait of Georgia over representative seasons and analyze metatranscriptomic profiles of the microorganisms present within intake water containing different pCO_2_ levels at a shellfish hatchery in British Columbia from June through October. Abiotic factors studied include pH, temperature, alkalinity, aragonite, calcite and pCO_2_. Community composition changes were observed to occur at broad taxonomic levels, and most notably to vary with temperature and pCO_2_. Functional gene expression profiles indicated a strong difference between early (June-July) and late summer (August-October) associated with viral activity. The taxonomic data suggests this could be due to the termination of cyanobacteria and phytoplankton blooms by viral lysis in the late season. Functional analysis indicated fewer differentially expressed transcripts associated with abiotic variables (e.g., pCO_2_) than with the temporal effect. Microbial composition and activity in these waters varies with both short-term effects observed alongside abiotic variation as well as long-term effects observed across seasons. The analysis of both taxonomy and functional gene expression simultaneously in the same samples by environmental RNA (eRNA metatranscriptomics) provided a more comprehensive view for monitoring water bodies than either would in isolation.

## Introduction

Metatranscriptomics characterizes expressed genes (i.e., RNA transcripts) that are present in an environmental sample. These transcripts may be within the cell, or in the extracellular space. While metagenomics profiles the taxonomy of a sample, metatranscriptomics can profile the biological functions that are present in the sample and are active or differentially accumulated in particular environments (Aguiar-Pulido et al. 2016). Metatranscriptomics can also be used to deduce taxonomic information for dominant taxa in communities (Shi et al. 2011; Neves et al. 2017; Marcelino et al. 2019), in particular with longer transcriptome contigs, which are expected to produce correct taxonomic assignments (Shakya et al. 2019). Metatranscriptomics is typically conducted through next-generation sequencing, which has the benefit of identifying novel genes and functions not known to be used by the identified taxa (Gilbert et al. 2008). Millions of marine microorganisms and viruses occur within a millilitre of seawater and their abundance and community composition varies with regions and environmental variables (Gilbert et al. 2011; Wigington et al. 2016; Finke et al. 2017; Yu et al. 2018; Gregory et al. 2019). A large scale global study found indications that the functional gene content in marine microbial samples is largely shaped by taxonomic composition (Salazar et al. 2019). However, the detailed distribution of generated community functions and their link to environmental variables remains to be studied (Moran 2015). Altogether, the environmental metatranscriptomic approach has strong potential for profiling active processes and community composition under changing conditions in marine molecular ecology (Moran et al. 2013; Cristescu 2019).

Coastal water microbial communities are shaped by widely varying environmental conditions, including temperature, salinity, and especially pCO_2_ and pH. (Salisbury et al. 2008; Joint et al. 2011; Ray et al. 2012; Lv et al. 2016; Lee et al. 2017). These are factors that are impacted by seasons, tides, or even biological activity. Microbes tolerate pH fluctuations under regular conditions and co-vary with water bodies in the short-term (Joint et al. 2011), but generally adjust in the long term. Microbial, bacterial and protist communities show significant responses to high pCO_2_ concentrations (Ray et al. 2012; Zhang et al. 2015; Thomson et al. 2016). In the past, average pCO_2_ concentrations have increased from 280 parts per million (ppm) (Friedlingstein et al. 2019) to currently 400 ppm (Blunden and Arndt 2020) and are deemed to increase to an average of 1000 ppm by the end of this century, with an associated rise of sea surface temperature by 1°C and drop of pH by 0.29 (Kirtman 2013; Pachauri 2014; Bindoff 2019).

Many microorganisms are expected to vary with temperature and/ or pCO_2_, and this may include shellfish pathogens such as *Vibrio* (Gomez-Leon et al. 2005; Green et al., 2019), *Roseovarius* (Maloy et al. 2007), and *Mikrocytos* (Carnegie et al. 2003), as well as harmful algae (Landsberg 2002) and viruses (Renault et al. 2014). By the nature of their intertidal habitat and filter feeding lifestyle, in particular oysters are affected by varying environmental conditions and microbiomes (Wegner et al. 2013; Lokmer et al. 2016;

Cho 2019). As a consequence, changes in microbial communities are expected to affect oyster hatcheries. Interestingly, for detecting an oyster parasite, environmental RNA (eRNA) was found to be superior to detect the species than when using environmental DNA (eDNA) (Merou et al. 2019). The impact of temperature increases and pCO_2_ on overall microbial communities and microbial activity is difficult to predict, but is expected to be significant with wide-ranging consequences. This serves as a motivation to develop comprehensive ways to improve our understanding of changes in marine microbial community composition and activity in response to environmental processes.

Here we compare the microbial composition, active genes, and enriched functional categories of differentially expressed genes within samples of intake water at a shellfish hatchery over a range of pCO_2_ levels, temperatures, and months. We apply comparative metatranscriptomics by first developing a *de novo* metatranscriptome assembly using the samples in the study. We take a taxonomic approach to view the community composition of each sample. In parallel, we quantify the relative expression levels of each transcript among the samples and perform differential gene expression analysis in relation to environmental metadata. Collectively, this work profiles hatchery intake water across 20 dates over a two-year period. Using environmental metadata, multivariate clustering, differential expression analysis, and functional enrichment analysis, we characterize these different sampling events in relation to date of collection and environmental parameters.

## Materials and Methods

### Water collection, RNA extraction and environmental variables

Water samples were collected from the ocean water intake at a Pacific oyster hatchery in Qualicum Bay, central East Coast Vancouver Island, British Columbia over a two year period, with the majority of samples collected in 2015 (Supplemental File S1). On each collection day, six litres of water were taken and filtered through sterile 0.22 μm cellulose acetate filters (Millipore, Burlington, MA). Filters were stored at −80°C. Temperature was measured by a mercury thermometer, salinity with a refractometer and pH with a glass probe pH meter (Jenco, San Diego, CA). Alkalinity was measured with a HI901 titrator (Hanna Instruments, Smithfield, RI). pCO_2_ was determined with a LI840A infrared gas analyzer (Li-Cor, Lincoln, NB). Aragonite and calcite were calculated with the CO2SYS.BAS program

(https://github.com/jamesorr/CO2SYS-Excel/blob/master/CO2sys_mod.bas). The filters were then used for RNA extraction by the Power Water RNA isolation kit (MoBio, Carlsbad, CA) following manufacturer’s instructions, including the alternate lysis step. Output total RNA was depleted for ribosomal RNA and prepped for RNA-seq using the Scriptseq Complete Gold (Epidemiology) kit (Illumina, San Diego, CA). A total of ~50 ng of ribosomal depleted RNA was used as an input to transcriptome libraries. Individual barcoded libraries were randomly multiplexed into groups of four samples using equimolar quantities and sequenced on a MiSeq using a v3 600 reagent kit (Illumina, San Diego, CA) to generate paired-end 250 bp reads. Environmental variables were analyzed in a principal component analysis (PCA) using the Vegan package (Oksanen et al. 2016) in the R environment (R Core Team, 2022), where missing values were imputed with unit averages.

### Bioinformatics and metatranscriptome assembly

Raw and quality trimmed sequence data were inspected with FastQC (Andrews 2010) and MultiQC (Ewels et al. 2016). Quality trimming was conducted to remove low quality reads and adapters with Cutadapt (Martin 2011) using flags -q 20 to remove < Q20 data from the 3’-end of the read and -m 50 to remove reads shorter than 50 bp. Results were output as an interleaved fastq. Putative ribosomal RNA (rRNA) reads were removed using SortMeRNA (Kopylova, Noé, and Touzet 2012) using all suggested rRNA databases, and therefore enriching for putative messenger RNA (mRNA) *in silico*.

Two different approaches were taken to assemble the reference metatranscriptome. First, a reference metatranscriptome was assembled using the metatranscriptome-optimized assembler IDBA-tran (Peng 2012) with a majority of the samples (16 / 20 samples; 191,303,324 reads), not including all samples due to computational constraints. Samples used for this approach comprised an equal number of samples from both ends of the pCO_2_ range present in the collections. This assembly was conducted with default settings and run on 56 threads. Second, a reference metatranscriptome was generated by first assembling each of the 20 libraries individually using IDBA-tran, and then merging these 20 assemblies into a single assembly using CD-HIT-EST (Li and Godzik 2006). CD-HIT-EST merged contigs at 95% similarity, and dedupe.sh of BBTools (Bushnell, Rood, and Singer 2017) was used to de-duplicate with default parameters (i.e., the ‘merged assembly approach’). These assemblies were compared based on the total number of contigs, total length, multi-mapping proportions, and mapping percentages to select the best assembly.

Reads for each sample were aligned against the reference metatranscriptome using Bowtie2 (Langmead and Salzberg 2012) in end-to-end mode allowing for multi-mappings. A maximum of 40 alignments were retained for each read. Alignments were then filtered to remove low quality mappings (i.e., retain mapq ≥ 2). Retained alignments were quantified using eXpress (Roberts et al. 2013). Effective counts from eXpress were output into a table in R, and imported into edgeR (Robinson and Oshlack 2010). Filtering was conducted to only retain contigs against which at least five reads mapped in the sample with the fewest reads (i.e., 3.86 counts per million; CPM), and requiring that the contig was represented at this CPM level or higher in at least five samples. Retained transcripts were normalized for library size using the TMM normalization method of edgeR v.3.28.1.

### Taxonomic community analysis

The expressed transcripts of the metatranscriptome were annotated for taxonomic identity using BLASTn (Camacho et al. 2009) against the nt database of NCBI, retaining a maximum of 100 alignments and descriptions per record. Best annotations were selected based on the e-value and a minimum cut-off at E < 10^-5^. Best match phylogenetic lineages of annotated transcripts were extracted with a custom python tool (see Data Accessibility) based on the subject sequence id using the ranked lineage database (NCBI), and exporting different levels of the taxonomy. The taxonomic overview and characterization used all expressed genes, at the kingdom, phylum and order level, and further analysis was conducted at the genus level. A canonical correlation analysis (CCA) of genus abundances and environmental variables, as well as ANOVA tests for significance of regressions were performed with the vegan package in R. To compare community composition at different lineage levels, pairwise distances were calculated using the Bray-Curtis dissimilarity measure in vegan, and Mantel tests were performed using the ade4 package (Dray and Dufour 2007). Linear models of genus abundance versus pCO_2_ concentration and t-tests for genus abundance between seasons were conducted in R. Significant regressions were defined by p < 0.05, adjusted p-values were determined using the Benjamin-Hochberg procedure.

### Differential expression analysis

To view detailed gene expression trends among samples, log2 CPM values were used for multidimensional scaling (MDS) plots. The plotMDS function of *limma* v.3.42.2 (Ritchie et al. 2015) was used to generate MDS plots using both the leading log2-fold-change as well as a PCA (gene.selection = common). Samples were grouped into binary groups for pCO_2_ (low/normal versus high) and season for differential expression analysis. High pCO_2_ was considered when the value was greater than 700 ppm, and low/normal was considered less than 700 ppm. Early summer was considered as June through July, and late summer was considered for August through October (see Table 1). Expression levels for each transcript were analyzed in a generalized linear model (i.e., glmFit and glmLRT) in edgeR to analyze the effect of pCO_2_ and the effect of early vs. late summer, and their interaction. Genes with pairwise p ≤ 0.05 after Benjamini-Hochberg multiple test correction were considered differentially expressed.

**Table 1.** Summary statistics of ocean chemistry variables. pCO_2_ was the target abiotic variable of the study, but the effect of the other variables was also considered in the context of metatranscriptome profiles. For calcite and aragonite, the saturation states are shown.

|  | Min | Max | Mean | Median | Std. dev. |
| --- | --- | --- | --- | --- | --- |
| pH | 7.56 | 8.35 | 7.8 | 7.78 | 0.22 |
| Temperature (°C) | 11.4 | 16.7 | 14.4 | 14 | 1.5 |
| Salinity (‰) | 28 | 29 | 28.2 | 28 | 0.41 |
| Alkalinity (μmol/kg) | 795.1 | 2634.2 | 1501.1 | 1400 | 465.1 |
| Aragonite (Ω) | 0.34 | 2.72 | 0.93 | 0.65 | 0.69 |
| Calcite (Ω) | 0.54 | 3.15 | 1.39 | 1.04 | 0.86 |
| pCO<sub>2</sub> (ppm) | 81.2 | 1059.9 | 588.3 | 606.6 | 289.9 |

To annotate transcripts with functions, expressed transcripts were assigned UniProt descriptions and identifiers by using BLASTx (Altschul 1997) to align contigs against the Swiss-Prot database (UniProt 2017) using the pipeline go_enrichment (Eric Normandeau, see Data Accessibility), with flags -- max_target_seqs 1 in outfmt 6 format, and only retaining hits with E < 10^-5^. The UniProt identifier was used as an input for Gene Ontology (GO) enrichment analysis in DAVID bioinformatics (Huang, Sherman, and Lempicki 2009), using differentially expressed lists compared against all expressed genes in the metatranscriptome for those transcripts annotated with UniProt identifiers.

## Results

### Sampling and environmental conditions

Intake water at the commercial oyster hatchery on east coast Vancouver Island, BC was sampled on four separate days in 2014 (June-August) and on 16 days in 2015 (June-November), with the total samples collected being seasonally balanced between the early summer (i.e., June-July, n = 11) and late summer (i.e., August-October, n = 9). Environmental variables were measured from the intake water, including pH, temperature, salinity, alkalinity, aragonite, calcite and pCO_2_, as shown in Table 1 (for complete data see Supplemental File S1).

The focal variable of this study, pCO_2_, ranged from 81-1060 ppm (average = 588 +/-290 ppm). Following IPCC assessments (Gattuso 2014) we classify samples into low pCO_2_ < 400 ppm (n = 6), medium pCO_2_ 400-700 ppm (n = 4), and high pCO_2_ > 700 ppm (n = 10) samples. A two dimensional PCA of environmental conditions shows variability among samples, separating them by low, medium and high pCO_2_ conditions (Figure 1). Samples S25 and S35, having concentrations of 397 and 386 ppm pCO_2_ cluster close to the medium pCO_2_ samples. Samples S22, S26 and S36 have almost identical environmental conditions. The PCA describes 77% of total variation in the first two principal components, with 52% explained by PC1 and 25% by PC2. The most influential variables in the PCA are pCO_2_ and pH, separating the samples by their pCO_2_ classes. The contributions of pCO_2_ and pH are inverse, as are salinity and temperature. Calcium carbonate (CaCO_3_), aragonite and calcite differ in their variation from the main axes described by pH/ pCO_2_ and salinity/ temperature. Notably, the grouping of samples by environmental variables does not display clustering by sampling months and the associated early vs. late summer classification (Figure 1).

**Figure 1.**
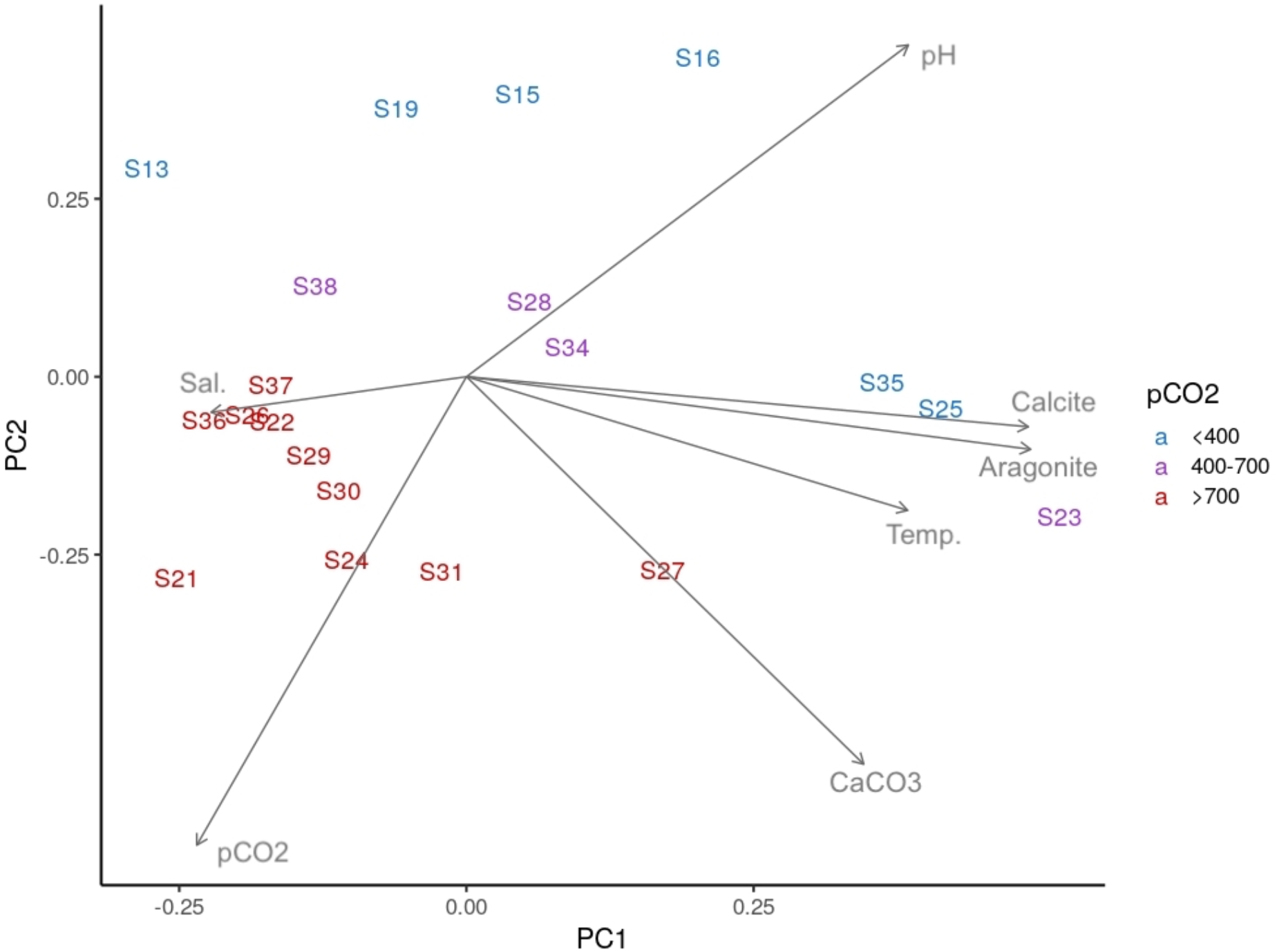
Principal Components Analysis (PCA) of samples based on environmental conditions. Samples are indicated as labels, their corresponding pCO_2_ classification is indicated by colour (blue=low, violet=medium, red=high). Arrow direction and length indicate the relative effect and strength of environmental variables. Here the positions of samples are entirely from environmental conditions, not based on taxonomic or gene expression data.

### Sequencing and assembly

The metatranscriptome libraries each yielded on average 14.7 M (s.d. = 2.8 M) paired-end reads. Depletion of rRNA in library preparation removed most of the rRNA from a majority of the samples, although for six of the 20 samples, 30-65% of sequenced library remained as rRNA (Figure S1). Residual rRNA was removed *in silico*, and is primarily comprised of bacterial 23s and eukaryotic 28s rRNA, but also archaeal rRNA.

The reference metatranscriptome was assembled from a total of 191,303,324 mRNA reads (33,050,961,278 bp) originating from pooling the non-rRNA data from 16 of the samples (n = 8 from each of low pCO_2_ and high pCO_2_). This input produced a final assembly of 8,003,896 contigs (total length = 2,468,090,451 bp; longest contig: 103,407 bp; N50 = 272 bp; number contigs > 500 bp = 600,691). This assembly was compared to other assemblies that used fewer libraries, or those that were individually assembled by sample then subsequently merged (i.e., ‘merged assembly’; see Methods). The collectively-assembled contigs with the highest number of input samples show fewer multi-mapping reads than did the merged assembly. The percentage of reads aligning a single time increases substantially until four libraries were added, then tapers off to not increase notably with eight, 12, or 16 libraries. The addition of more libraries after four libraries did not substantially increase the percentage of reads mapping, but it also did not increase redundancy as evaluated by multi-mapping. The 16 library collectively-assembled assembly also has a similar number of contigs and total length to the other assemblies (see Supplemental Results; Figure S2). Therefore, this collectively-assembled assembly was chosen to be used for all downstream functional analyses (i.e., the ‘final reference metatranscriptome assembly’).

Aligning reads from individual samples against the final reference metatranscriptome assembly resulted in an average alignment rate per sample of 56% (median = 57%; min. = 37%; max = 68%), with an average of 40% of the total alignments per sample with both read pairs aligning concordantly a single time. On average per sample, 12% of reads align concordantly more than once, which may indicate remaining redundancy in the assembly. Applying low expression filters removed the majority of contigs from the metatranscriptome, retaining 32,866 contigs with CPM ≥ 3.86 in at least 5 of 20 samples.

### Taxonomic analysis

To infer the community compositions of samples, the taxonomic lineages for expressed transcripts were analyzed. Taxonomic assignment was successful for 19,315 of the 32,866 expressed transcripts (59%), but the taxonomic resolution varied among contigs. For example, 89% of the annotated transcripts resolve to at least the phylum level, 80% to class, 79% to genus and only 30% to the species level. The data includes some expected non-microbial taxa such as fish or molluscs and some suspected false taxonomic assignments, but is vastly dominated by microbes. A total of 708 genera in 104 classes and 58 phyla are annotated, but only 10 microbial phyla are present at an abundance level of greater than 0.5% of the total reads. When combined, these 10 phyla account for over 95% of all reads, yet several other phyla are represented in the data as well (Supplemental File S2). These 10 dominant phyla show an abundance of reads assigned to bacteria (~74%), specifically *Proteobacteria, Bacteriodetes, Firmicutes, Actinobacteria* and *Cyanobacteria*. Overall, fewer reads are assigned to archaea (~11%), viruses (~9%) and eukaryotes (~1.6%). *Archaea* are represented by *Thaumarcheota* and *Euryarcheota*. Viruses are largely in the class *Caudovirales* (*Uroviricota*) which includes, for example, bacteriophages and cyanophages, and the *Nucleocytoviricota* that represent the nucleocytoplasmic large DNA viruses (NCLDV). Eukaryotes are in the phylum *Bacillariophyta*. Figure 2 shows the variation in community composition of these ten phyla among samples, *Proteobacteria* and *Bacteroidetes* are clearly dominating in most samples, especially in the early season. *Cyanobacteria* and *Bacillariophyta* are also mostly present in early season samples. In late season there is an increase in relative abundance for the *Thaumarchaeota* and *Euryarchaeota*, but especially the *Uroviricota* and *Nucleocytoviricota* viruses.

**Figure 2.**
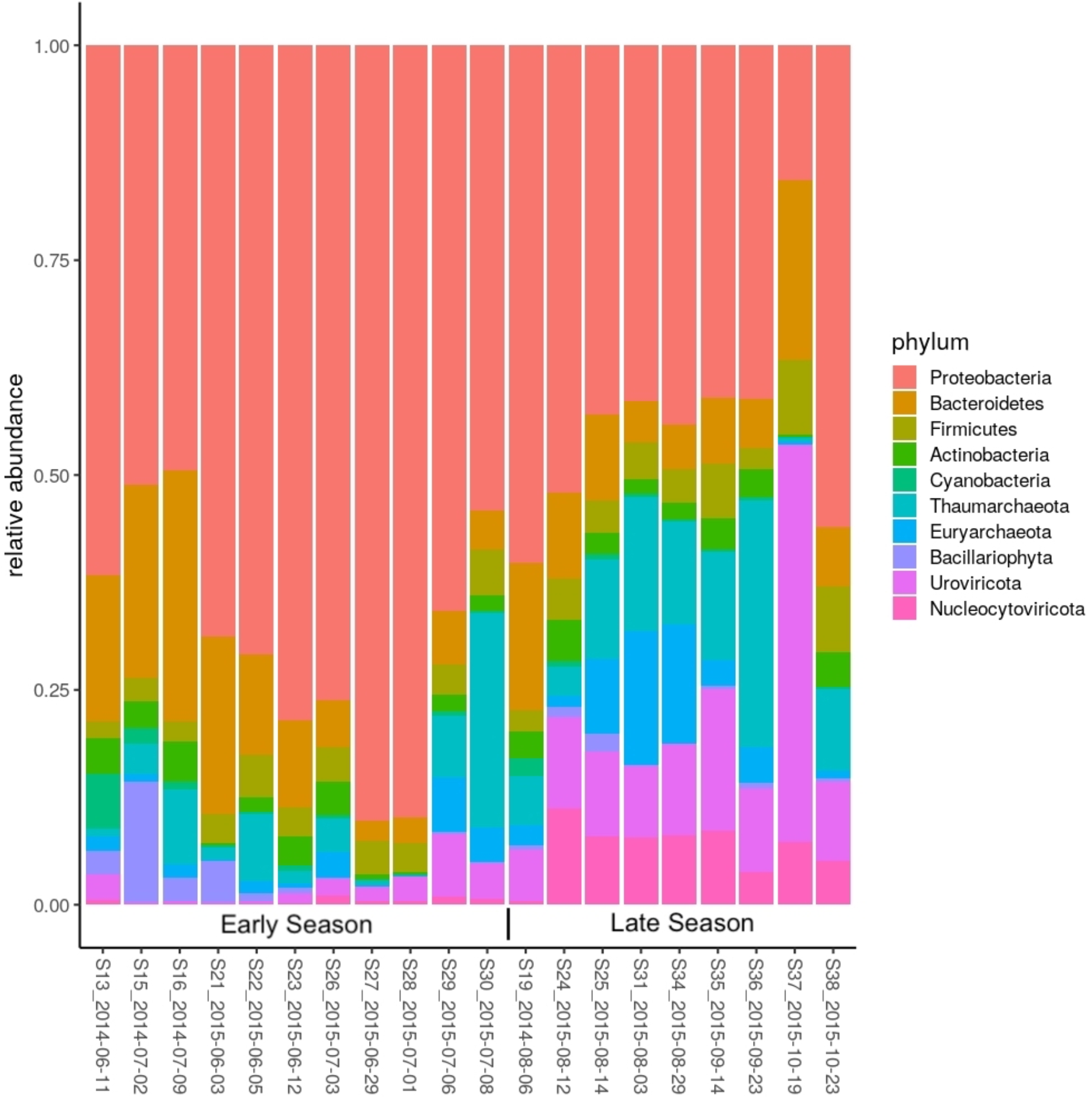
Community composition of the top 10 phyla across samples. Stacked bar-plot of the relative abundance (total number of alignments to taxa) across samples for microbial phyla above 0.5% total abundance, phyla are indicated by color coding in legend, samples are arranged by early season vs. late season.

The overall variation in community composition among samples was evaluated based on Bray-Curtis similarity in a pairwise distance analysis. When compared at different taxonomic levels, variations in community compositions are congruent, and this is true for comparisons of genus to class (R=0.97), class to phylum (R=0.91) and genus to phylum (R=0.90), all showing significant congruence (Mantel test p ≤ 0.01). A CCA of taxa composition for the 708 genera and environmental variables produces a significant model (P=0.013), explaining 56% of the total variation in taxon composition through environmental variables (Figure 3). Of that variation the first dimension (CCA1) describes 39% of the variation and the second dimension (CCA2) describes 27%. The relationship between sample composition as indicated by the sample labels and the genera (dots) are shown, where taxa are coloured according to the corresponding four kingdoms. Generally, kingdoms are distributed across the CCA, but eukaryotes show some grouping with samples S21, S15 and S13. Viruses show grouping with late summer samples S31, S34, S36 and S37. The effect and strength of environmental variables on the sample composition is indicated by vectors. Temperature and salinity, and pH and pCO_2_ describe the predominant dimensions of variation. A stepwise regression determined that temperature (p=0.001) and pCO_2_ (p=0.028) are significant environmental variables affecting the composition of samples. Additionally, early vs. late summer is significant (p=0.023) in separating samples by composition. Correlating all 708 genera to pCO_2_, our main environmental variable of interest, revealed 67 genera showing a linear correlation of their log_10_ transformed abundances to pCO_2_ concentrations (p < 0.05; Supplemental File S3), although of these, only two are significant based on multiple test corrected values (i.e., *Methyloceanibacter* and *Deferribacter*). The genera with the highest linear correlations are predominantly in the *Proteobacteria, Cyanobacteria*, and *Picornavirales* and *Caudovirales*. Similarly, 93 genera show significant variation between the early and late season samples, especially *Caudovirales* and *Algavirales* (Supplemental File S4). Of these, 23 are significant based on multiple test corrected values, including *Caudovirales*. The top ten genera for both models are summarized in Table 2.

**Figure 3.**
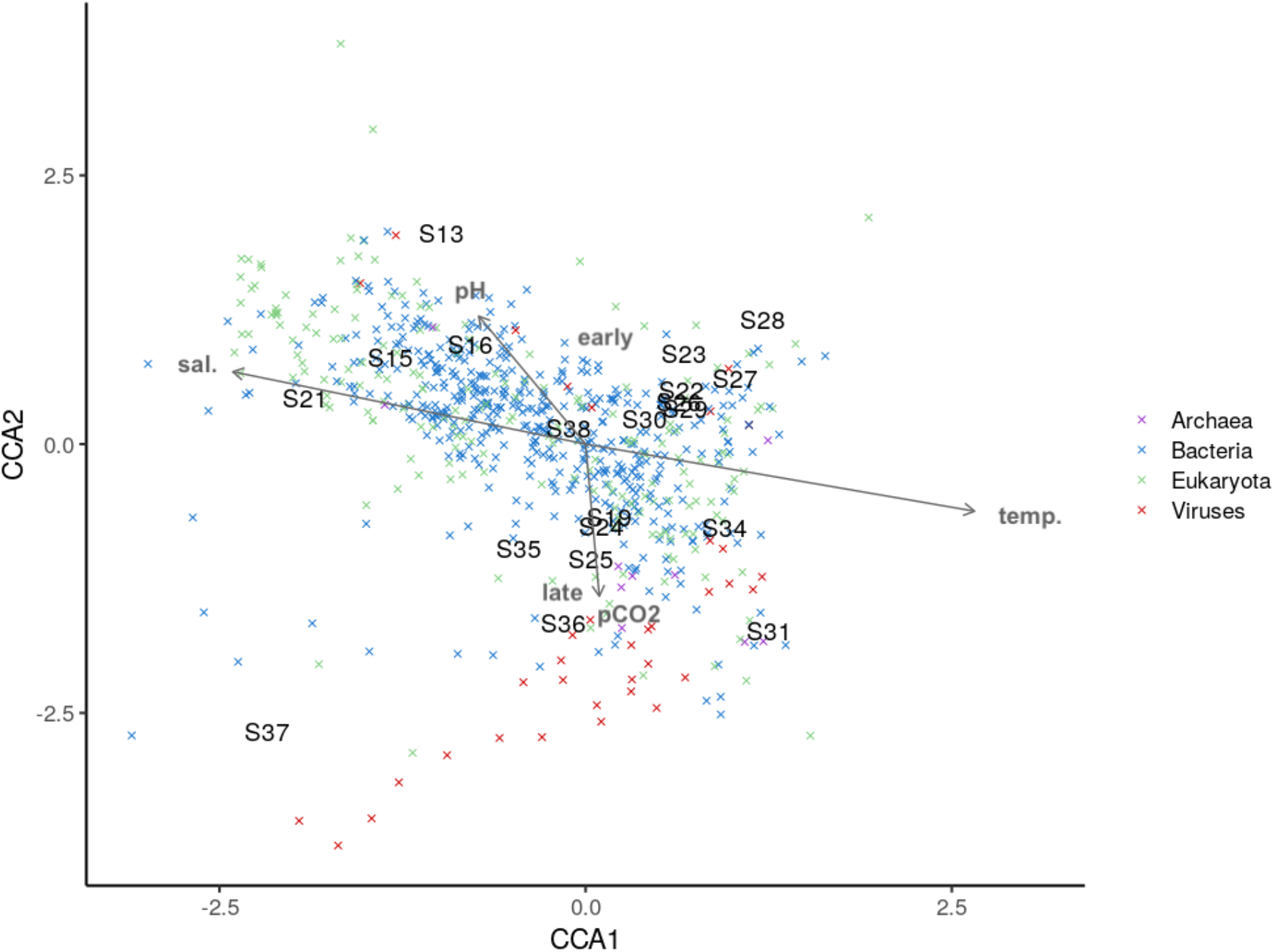
Canonical correspondence analysis (CCA) showing the effect of environmental variables on the variation in relative abundance of genera among samples. Samples are shown as labels and the relative association of genera is shown as crosses. Crosses for genera are color coded by their corresponding superkingdom. The effect of environmental variables on the composition of genera in the samples is indicated by grey arrows, the arrow length corresponds to scaled effect strength. The effect of early and late summer is indicated in grey labels.

**Table 2.** Top ten significant genera of linear models against low, medium, and high pCO_2_ concentration and T-test of early vs. late summer. P-values express the model significance and P-adj. is adjusted p-value with Benjamini-Hochberg. Statistic values indicate positive and negative correlations.

| **Linear model genus vs. pCO<sub>2</sub>** |  |  |  |  |  |  |  |
| --- | --- | --- | --- | --- | --- | --- | --- |
| R2 | P-value | P-adj | Slope | Genus | Order | Phylum | Superkingdom |
| 0.718 | <0.001 | 0.038 | 0.001 | *Methyloceanibacter* | *Rhizobiales* | *Proteobacteria* | *Bacteria* |
| 0.597 | <0.001 | 0.038 | -0.002 | *Deferribacter* | *Deferribacterales* | *Deferribacteres* | *Bacteria* |
| 0.552 | <0.001 | 0.063 | -0.001 | *Gossypium* | *Malvales* | *Streptophyta* | *Eukaryota* |
| 0.614 | 0.001 | 0.098 | -0.002 | *Thioalkalivibrio* | *Chromatiales* | *Proteobacteria* | *Bacteria* |
| 0.694 | 0.001 | 0.102 | -0.002 | *Mitrocomella* | *Leptothecata* | *Cnidaria* | *Eukaryota* |
| 0.466 | 0.001 | 0.102 | -0.002 | *Sogarnavirus* | *Picornavirales* | *Pisuviricota* | *Viruses* |
| 0.460 | 0.001 | 0.102 | -0.001 | *Methylovorus* | *Nitrosomonadales* | *Proteobacteria* | *Bacteria* |
| 0.447 | 0.001 | 0.108 | -0.001 | *Methylophilus* | *Nitrosomonadales* | *Proteobacteria* | *Bacteria* |
| 0.462 | 0.001 | 0.108 | -0.001 | *Aeromonas* | *Aeromonadales* | *Proteobacteria* | *Bacteria* |
| 0.470 | 0.002 | 0.155 | 0.001 | *Steinhofvirus* | *Caudovirales* | *Uroviricota* | *Viruses* |
| **T-test genus vs. early and late summer** |  |  |  |  |  |  |  |
| Statistic | P-value | P-adj | Stderr | Genus | Order | Phylum | Superkingdom |
| 7.053 | <0.001 | 0.001 | 0.121 | *Coregonus* | *Salmoniformes* | *Chordata* | *Eukaryota* |
| 6.884 | <0.001 | 0.002 | 0.118 | *Biomphalaria* |  | *Mollusca* | *Eukaryota* |
| 4.129 | 0.001 | 0.030 | 0.169 | *Leucotheavirus* | *Caudovirales* | *Uroviricota* | *Viruses* |
| 3.759 | 0.004 | 0.088 | 0.207 | *Nesterenkonia* | *Micrococcales* | *Actinobacteria* | *Bacteria* |
| 3.713 | 0.002 | 0.051 | 0.217 | *Glaciecola* | *Alteromonadales* | *Proteobacteria* | *Bacteria* |
| 3.693 | 0.002 | 0.062 | 0.164 | *Emcibacter* | *Emcibacterales* | *Proteobacteria* | *Bacteria* |
| 3.614 | 0.002 | 0.062 | 0.125 | *Glaesserella* | *Pasteurellales* | *Proteobacteria* | *Bacteria* |
| 3.566 | 0.004 | 0.095 | 0.217 | *Nephromyces* | *Nephromycida* | *Apicomplexa* | *Eukaryota* |
| 3.560 | 0.006 | 0.110 | 0.194 | *Tannerella* | *Bacteroidales* | *Bacteroidetes* | *Bacteria* |
| 3.221 | 0.006 | 0.110 | 0.112 | *Borrelia* | *Spirochaetales* | *Spirochaetes* | *Bacteria* |

### Gene expression analysis

After the low expression filter was applied, in total 22,121 (67%) of the expressed contigs (n = 32,866) were functionally annotated with UniProt identifiers (BLASTx E < 10-5). All gene expression analysis was conducted using both the annotated and unknown transcripts except for functional enrichment analysis, which depends on the UniProt identifier.

Using the filtered expression data, an unsupervised multidimensional scaling plot (MDS plot) was used to group samples by similarity in gene expression (Figure 4). Based on gene expression signatures this plot indicates similarity among samples from a common season, irrespective of sampling year, where samples from June-July (early summer) are separated from August-October (late summer/fall). Early season samples (blue) and late season samples (red) separated on PC1 (Figure 4), with one sample being an exception, S19, which groups outside of its season. Notably, S19 was the only sample from 2014 that was collected in August or later (collected Aug. 6^Th^, 2014; Supplemental File S1). No grouping within the MDS plot was observed for an effect of year nor of technical aspects of sample handling.

**Figure 4.**
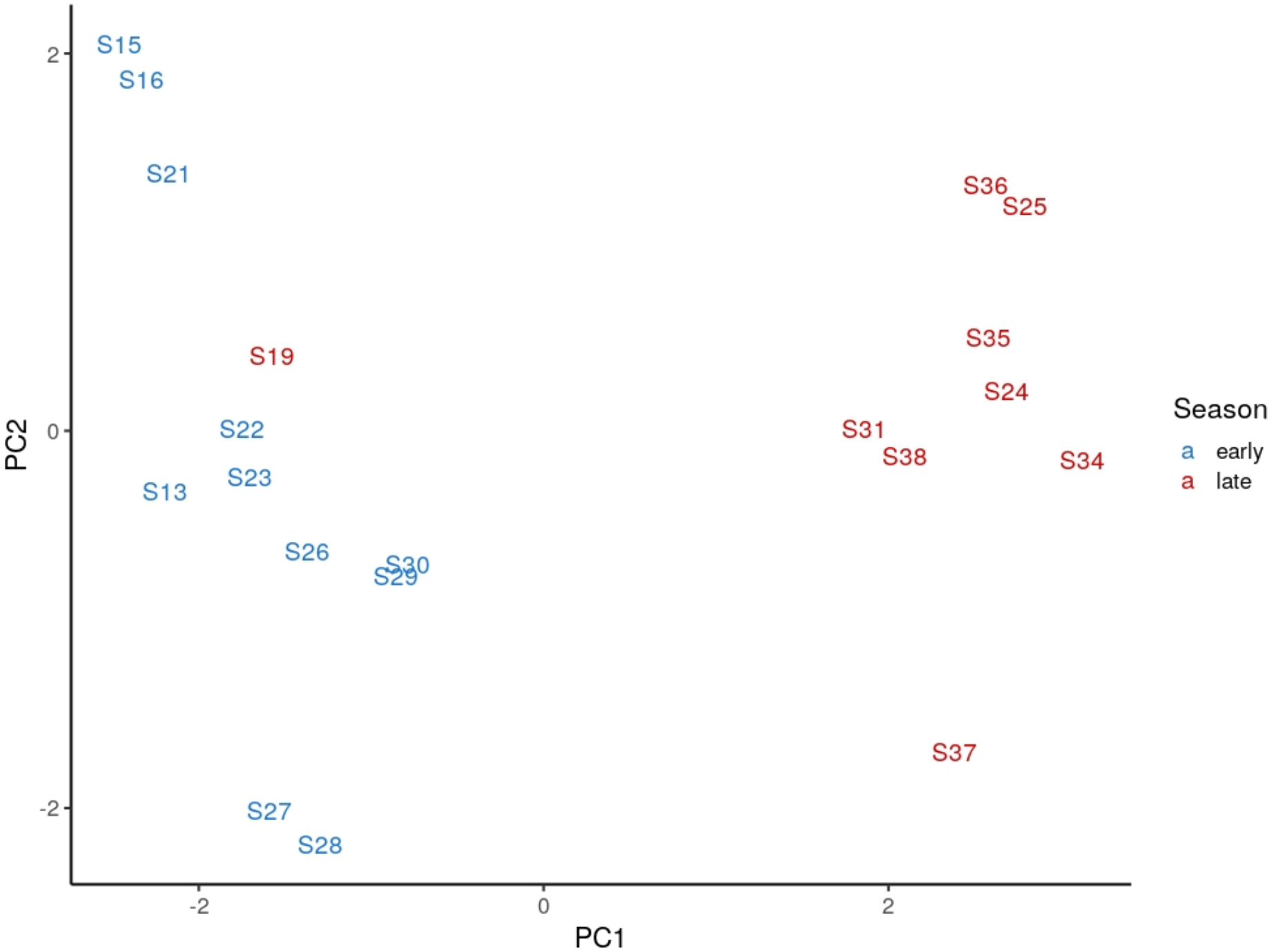
Unsupervised multidimensional scaling (MDS) plot on samples based on gene expression. Dimension 1 explains the most variation, separating the late and early season samples. Samples are labeled by sample number and colors indicate early vs. late season. Samples S13, S15, S16, and S19 are from 2014, and the rest are from 2015. Full details on samples can be viewed in Supplemental File S1.

Based on the observations of the effect of pCO_2_ as a key environmental variable on taxonomic composition and a focal variable of the study, a differential expression analysis was conducted using pCO_2_ separated into either high or medium/low levels, and early summer vs. late summer as binary explanatory variables, as well as their interaction. There is a larger effect of early vs. late summer than of pCO_2_, where 2,765 transcripts are found differentially expressed based on early vs. late, and 720 based on pCO_2_. Of these, 45 are differentially expressed in both contrasts (Supplemental File S5). Of the transcripts affected by season, 318 are over-expressed in early summer, and 2447 are over-expressed in late summer/early fall. Of the transcripts affected by pCO_2_, 553 are over-expressed by high pCO_2_ and 167 are under-expressed. There are no transcripts showing a significant interaction effect of sampling period and pCO_2_.

Differentially expressed transcripts were used for Gene Ontology (GO) enrichment analysis (Table 3). Annotated transcripts overexpressed in the early summer (n = 318) are enriched for metabolic processes and biosynthesis (e.g., cellular macromolecule metabolic process; n = 32 genes; p = 0.01), and those overexpressed in the late summer are most notably enriched with viral process (biological process; n = 53; p << 0.0001) and virion (cellular component; n = 30; p << 0.0001). The viral process category is mainly enriched with transcripts annotated from phage taxa (n = 40 of 53 transcripts; Supplemental File S6).

**Table 3.** Gene Ontology (GO) enrichment for pCO_2_ and season. GO enrichment analysis indicates the effect of pCO_2_ and season on several biological processes, most notably a highly significant enrichment for viral processes in the late season. Columns shown are the GO Term, count in gene list, p-value for enrichment test, count in background list, and fold enrichment. Full GO enrichment results are presented in Supplemental Materials.

|  | GO Term | Count | P-value | Count<br>(background) | Fold<br>Enrich. |
| --- | --- | --- | --- | --- | --- |
| **Overexpr. in**<br>**high pCO<sub>2</sub>** | GO:0009236~cobalamin<br>biosynthetic process | 5 | 0.0030 | 18 | 7.9 |
|  | GO:0006260~DNA replication | 17 | 0.0048 | 226 | 2.1 |
|  | GO:0019058~viral life cycle | 8 | 0.015 | 75 | 3.0 |
|  | GO:0039693~viral DNA<br>genome replication | 4 | 0.020 | 17 | 6.7 |
| **Overexpr. in**<br>**low pCO<sub>2</sub>** | GO:0051604~protein<br>maturation | 6 | 6.1E-06 | 30 | 21.0 |
|  | GO:0006950~response to<br>stress | 10 | 0.0032 | 344 | 3.1 |
|  | GO:0006457~protein folding | 5 | 0.0067 | 83 | 6.3 |
| **Overexpr. in**<br>**early season** | GO:0003735~structural<br>constituent of ribosome | 8 | 0.0059 | 134 | 3.6 |
| **Overexpr. in**<br>**late season** | GO:0016032~viral process | 53 | 7.3E-42 | 91 | 9.7 |
|  | GO:0006260~DNA replication | 55 | 1.8E-20 | 226 | 4.1 |
|  | GO:0051701~interaction with<br>host | 23 | 5.1E-16 | 45 | 8.5 |
|  | GO:0046718~viral entry into<br>host cell | 17 | 1.2E-13 | 27 | 10.5 |

Genes overexpressed in high pCO_2_ include cobalamin biosynthetic process (n = 5, p = 0.003), organic substance biosynthetic process (n = 81, p = 0.004), and DNA replication (n = 17, p =0.005). Genes with lower expression at high pCO_2_ include protein maturation (n = 6, p << 0.001) and response to abiotic stimulus (n = 6, p < 0.001).

## Discussion

### Profiled characteristic conditions indicate short-term environmental fluctuations

The collected samples reflect typical intake water of oyster hatcheries from standard aquaculture practices on East Coast Vancouver Island (Helm 2004), spanning over several months and two years. The variables were analyzed in a PCA to understand the covariation of environmental variables and how they shape the sampling conditions. The variation in environmental data describes the characteristic water flow and carbon chemistry of intertidal oyster habitats and the region of sampling (Strait of Georgia) (Dickinson et al. 2012; Ianson et al. 2016); salinity and temperature are inversely affected by influxes of cold seawater with relatively high salinity and warmer fresh water with lower salinities. Water with higher pCO_2_ concentrations is characterized by lower pH values and the combined effect of pH, pCO_2_, and salinity is reflected in the CaCO_3_ concentrations, with calcium being one of the salts in seawater. This relationship shows in the sample separation by low, medium and high pCO_2_ values and the force loadings of environmental variables. Calcite and aragonite are both derivatives of CaCO_3_, varying based on the pCO_2_ and the pH of water (Doney 2010), and their influence is thus mediated between these two variables. The clustering of samples observed when considering environmental data highlights the impact of pCO_2_ and salinity and suggests that variations in environmental conditions observed in the scale of the present study (two years) are driven by short term influxes of fresh or marine waters through tidal cycles and upwelling of deep waters.

### Taxonomic composition varies with abiotic factors and early and late summer

The microbial community analysis here is based on sequences with taxonomic assignment. However, we expect this to be a representative description, with the possible exception of largely uncharacterized taxa in reference databases. Derived from taxonomic assignments of contigs and their respective expression levels, community similarity among samples proved to be congruent across different taxonomic levels. This validates the approach of using taxonomic assignments down to the genus level. Importantly, it also indicates that shifts in community composition happen at higher taxonomic levels and not just at the genus or species level. Contigs assigned to non-microbial taxa or terrestrial taxa are expected to be either the product of dispersed tissue or cells in the water column or alternatively incorrect taxonomic assignment.

Across samples the microbial communities are clearly dominated by *Proteobacteria, Bacteriodetes, Firmicutes*, *Actinobacteria* and *Cyanobacteria*. Both eukaryote and archaea microorganisms are notably less represented, and even matched in abundance by viral sequences. The described bacterial phyla are commonly found to be dominant in marine (Sunagawa et al. 2015) and coastal waters (Yung et al. 2016; Yu et al. 2018), and constitute the microbiome of oysters themselves (Trabal et al. 2012; Lokmer et al. 2015; Lokmer et al. 2016; Dubé et al. 2019; Stevick et al. 2019). In particular, *Proteobacteria (Vibrionales*) and *Bacteriodetes (Flavobacteria*) represent common marine microbes and pathogens (Gomez-Leon et al. 2005; Schulze et al. 2006; Paillard et al. 2008; Chen et al. 2017). Observations by Stevick and co-workers (Stevick et al. 2019) found that *Cyanobacteria* were among the dominant phyla in the oyster rearing water communities and *Synechococcus* is a ubiquitous cyanobacterial genus in coastal environments (Partensky et al. 1999; Tai and Palenik 2009). Similarly, the abundant eukaryote *Bacillariophyta* and *Chlorophyta*,both present in the samples taken here, are common phytoplankton in coastal waters (Worden, Nolan, and Palenik 2004; Armbrust 2009).

Matching the microbial community, the dominant *Caudovirales* includes general bacteriophages and specifically cyanophages (Weinbauer and Rassoulzadegan 2004). The present results are thus mirroring the general ubiquity of heterotrophic bacteria, but also indicating their lysis. The *Nucleocytoviricota* include giant viruses commonly infecting protists (Fischer et al. 2010; VanEtten et al. 2010), but also phycodnaviruses infecting eukaryote phytoplankton (e.g., *Chlorophyta*), both being abundantly present in coastal waters (VanEtten et al. 1982, 2002). Taken together, the observation of a low presence of cyanobacteria sequences and *Bacillariophyta* sequences alongside high abundance of plankton viruses and phages in some late season samples, may indicate the lysis of cyanobacteria and algae blooms later in the season. This observation would match phytoplankton bloom patterns described for the northern Strait of Georgia where diatoms (*Bacillariophyta*) and prasinophytes (*Chlorophyta*) are mostly dominant with only periodic cyanobacteria blooms (Del Bel Belluz et al. 2021). Additionally, the general decrease in *Proteobacteria* in the late season samples is observed alongside increased presence of phage (*Uroviricota*)activity, which also fits with the understanding that bacterial blooms are terminated by viral lysis. As well, viruses show a correlation to late summer samples in the canonical correspondence analysis and the t-test following multiple test correction (i.e., *Caudovirales*).

In the linear models of taxa to pCO_2_, our main environmental variable of interest, 67 correlations are significant based on individual tests (p < 0.05), but only two are significant based on multiple test corrected p-values, and so the 65 taxa should be considered as putatively characteristic of changes in pCO_2_ due to water influx. Future studies may indicate the broader reliability of the finding in these taxa. Several of these 67 taxa, including *Proteobacteria, Bacteroidetes, Cyanobacteria, Firmicutes* and *Deferribacteres* match bacterial taxa with abundances that are associated with tidal cycles and salinity (Lee et al. 2017; Chen et al. 2019). There remains a large portion of unexplained variation in the taxonomic data when all taxa are considered. The CCA shows that only about 50% of the variation among microbial communities could be explained by the combined environmental variables. The effect of environmental variables on the taxonomic community data in the CCA matches their interplay in the PCA based on environmental data (Figure 1). The CCA also confirms that early vs. late summer sampling time has a significant effect on the community composition, most notably for viruses, although overall to a lesser effect than observed in the functional gene expression results (see below). We could not identify the abiotic variables responsible for the early vs. late summer effect, which indicates other variables that may be involved in community differences are missing (e.g., nutrients or irradiance levels, among others).

In any case, the observed effect and explanatory levels of environmental variables matches previous studies, where temperature has been shown to be a main variable correlated to community composition of coastal marine bacterioplankton (El-Swais et al. 2015; Yung et al. 2016; Yu et al. 2018). Further, Yu and co-workers also established the effect of pH that corresponds to our observations on the influence of pCO_2_ on community composition. Therefore, although some abiotic or biotic variables may be missing from the study, the results fit with other studies on abiotic factors influencing microbial communities.

Overall, the data show that the microbial communities in coastal waters vary at high taxonomic levels with season. The communities further vary alongside with environmental variables that are driven by tidal cycles and upwelling. As demonstrated, metatranscriptome data can be used to monitor the presence of a wide range of microbes and putative pathogens in seawater, expanding on the use of specific microbe probing (e.g., Merou et al. 2019).

### Differential RNA transcript abundance reveals functional variation between early and late summer

Environmental variables are not only expected to alter microbial community composition, but also to influence the function of cells on the transcriptomic level. Further, a change in community composition does not necessarily indicate a change in functions within the community if one taxon is replaced by a similar taxon with similar functions. A study comparing taxonomic and functional composition in marine microbial samples revealed a disconnect between taxonomy and functions (Louca et al. 2016). In the functional transcriptome analysis of the present study, early and late summer sampling shows a clear effect on overall transcriptome expression, but environmental variables including pCO_2_ do not show clear groupings of samples in the gene expression data. Consequently, similar to the community composition results, here functional variation is expected to have been influenced by factors outside of the measured environmental variables (e.g., nutrients and irradiance levels, among others). Observing this pattern in gene expression despite having samples spread out over two years suggests some consistency to this seasonal trend. However, the grouping of the only August sample (S19) from 2014 not with the late summer but rather with the early summer unlike the August samples in 2015 may indicate a difference in the exact timing of the putatively seasonal shift in expression. Without more samples from 2014 after the August sample (S19), it is not possible to tell if the late season signature would appear again, but the early samples from both years consistently group. To determine if this late season signature persists in multiple years, and when the switch from early to late occurs each year, would require future work.

The early vs. late summer effect is most notably enriched for transcripts involved in phage viral activity, matching the findings in the taxonomic analysis. Viruses are recognized as major components in marine ecosystems (Suttle, 2007; Brussaard et al. 2008), but their replication dynamics remain elusive. However, in the North Pacific Ocean it has been observed that viral productivity and abundance is higher in summer (July) than in winter (Jan-Feb) (Gainer et al. 2017), leading these researchers to conclude that seasonality is an important consideration to understand viral dynamics (Jiang and Paul 1994; Tsai et al. 2013). A study within a Korean bay over different seasons found that the number of reads and unique species of viruses identified differed depending on month; the most viral reads were found in March and December and the fewest reads in June and September (Hwang 2017). It is recognized that viral composition changes depend seasonally on a range of factors including temperature, salinity, dissolved oxygen, primary production and nutrient concentrations (Brum et al. 2015; Fuhrman et al. 2015). Generally, viral abundance, specifically for dsDNA viruses, has been found to correlate with nutrient concentration, as well as heterotrophic bacteria abundance (Wigington et al. 2016; Finke et al. 2017). Viruses infecting bacteria (i.e., bacteriophages) are typically considered among the dominant group of viruses in marine environments (Breitbart et al. 2002; Steward et al. 2013). In the Korean bay study, 73% of the viral reads were from bacteriophage, and 26% were from algal viruses, with only 1% involving other viruses (Hwang et al. 2017). The effect of pCO_2_ was also investigated in the differential expression analysis, but this had a lesser effect than that the temporal effect observed. Interestingly, in the present study, overexpression of protein chaperones was observed at low pCO_2_, which is the opposite to previous observations where metatranscriptomic responses of overexpressed chaperonin transcripts was observed in high pCO_2_ mesocosms (Gilbert et al. 2008).

Marine microbial communities are often comprised of a few dominant species and many rare ones, and results from the Tara Oceans Expedition found approximately 37 k bacterial and archaeal species, 100 k protist groups and 5.5 k double-stranded bacterial and archaeal virus populations (summarized in Moran 2015). The present study is successful in assessing this high degree of diversity in gene expression profiles among samples. As is suggested from the community composition and transcript activity of the present study, previous metatranscriptome studies have also found a substantial increase in viral transcripts after a phytoplankton bloom, presumably due to infected cells in lytic stage (Gilbert et al. 2008).

Overall, taxonomic and transcriptomic profiling complemented each other and when combined provided a comprehensive view of the changes observed in this study. This matches the observation by Salazar et al. (2019) that the correlation between taxonomic composition and functional composition is variable. The community composition analysis provided more information on taxa associated with abiotic factors, and the functional analysis highlighted the large effect of viral activity that differed over time during the summer months. Being able to assess viral activity alongside microbial data is a major advantage of the metatranscriptomic (eRNA) approach applied here to assess marine microbial communities.

## Conclusions

In the present study, microbial communities were derived from metatranscriptome data, which avoided primer bias that can occur in amplicon-based approaches, and captures data from all superkingdoms. The observed variability among microbial communities was found to be associated with both temperature and pCO_2_, as well as other changes that occurred between early and late summer. Changing environmental conditions may occur over short time scales through upwelling or currents, or may have large, structured changes that occur temporally. These temporal and abiotic factors appeared to be disconnected and were two different trends occurring in the marine environment. The functional gene expression analysis pointed to a strong difference in viral activity moving into the late season, and a much lesser effect of abiotic factors such as pCO_2_, temperature and salinity. Together, these analyses provide community composition and functional gene differences associated with abiotic factors and time, and likely captured viral termination of bacterial, cyanobacterial and algal blooms later in the season. Being able to indiscriminately assess eukaryote, prokaryote and viral taxa, describing a combined bloom and viral lysis across several host virus systems proved to be the advantage of using metatranscriptome analysis. The improved understanding of changes in marine microbial communities with abiotic factors helps to monitor water bodies. Metatranscriptomics of eRNA allowed the characterization of both community and gene expression activity changes simultaneously, providing a comprehensive view of these dynamic water bodies.

## Acknowledgements

This project would not have been possible without the involvement, contributions, and sample provision from the industry partner, Island Scallops (ISL; CEO Robert Saunders). J. Finke and B. Sutherland were supported during this work by a grant from the Gordon and Betty Moore Foundation (GBMF, Grant number 5600) awarded to C. Suttle and K. Miller. This project was supported by an Aquaculture Collaborative Research and Development Program (ACRDP) grant from Fisheries and Oceans, Canada, awarded to K. Miller (Grant number P-14-02-001). The authors declare that they do not have any conflicts of interest in this work.

## Data Accessibility

Supplemental files including the reference metatranscriptome (fasta), quantified transcript expression levels (.csv), and annotation for the functional and taxonomic analyses (.txt) are available on FigShare: https://doi.org/10.6084/m9.figshare.18128966.v1

Pipeline for analyzing metatranscriptome data: https://github.com/bensutherland/eRNA_taxo Pipeline used for annotating transcripts: https://github.com/enormandeau/go_enrichment Script to extract taxonomic lineages: https://github.com/janfelix/bioinformatic_tools/blob/master/taxid2lineage.py

## Supplementary Materials

**Supplemental Figure 1.**
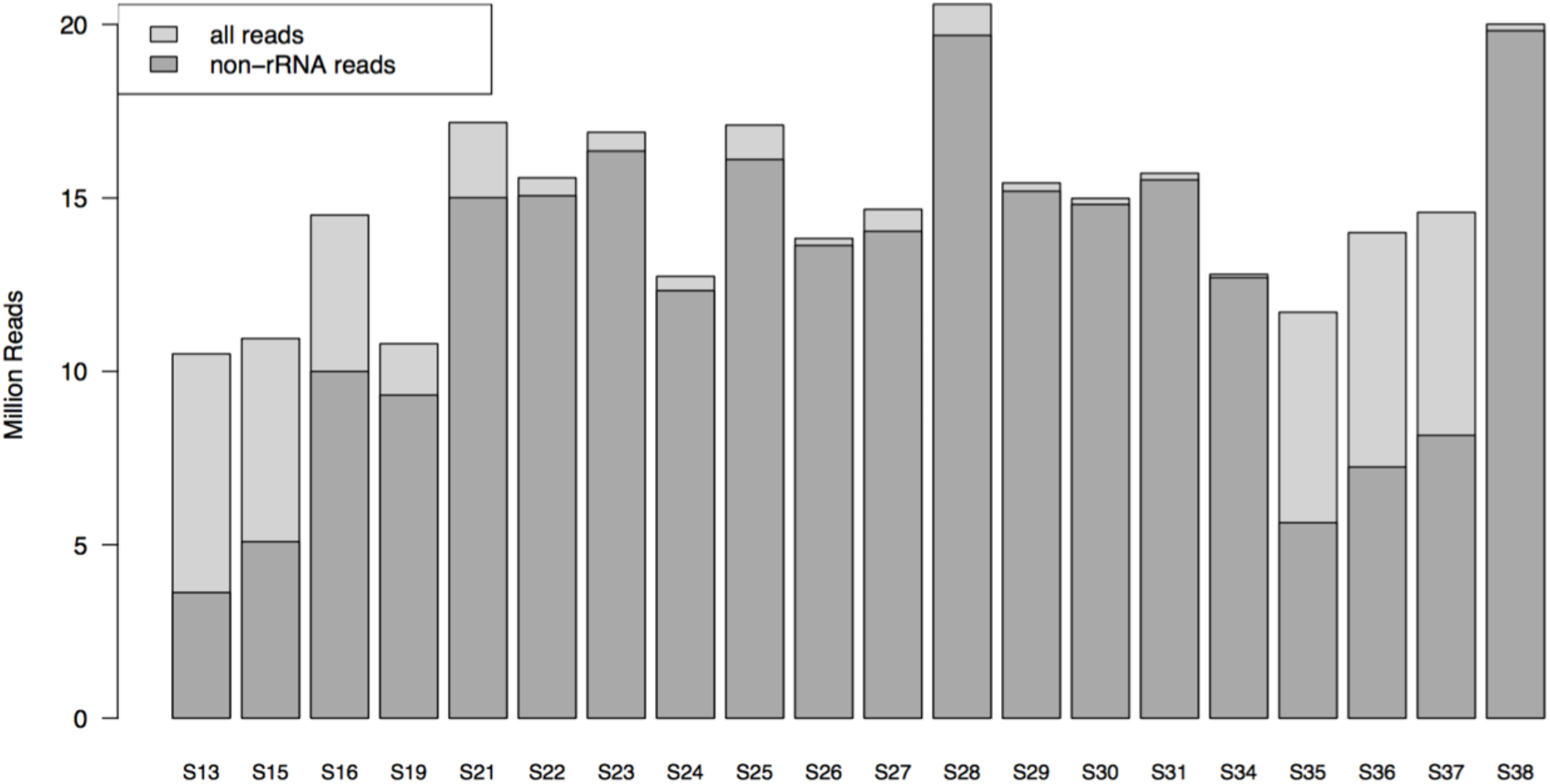
Proportion of non-rRNA and rRNA reads per sample. Results indicate that for a majority of the samples, the rRNA depletion was successful, enriching the amount of messenger RNA for analysis.

**Supplemental Figure 2.**
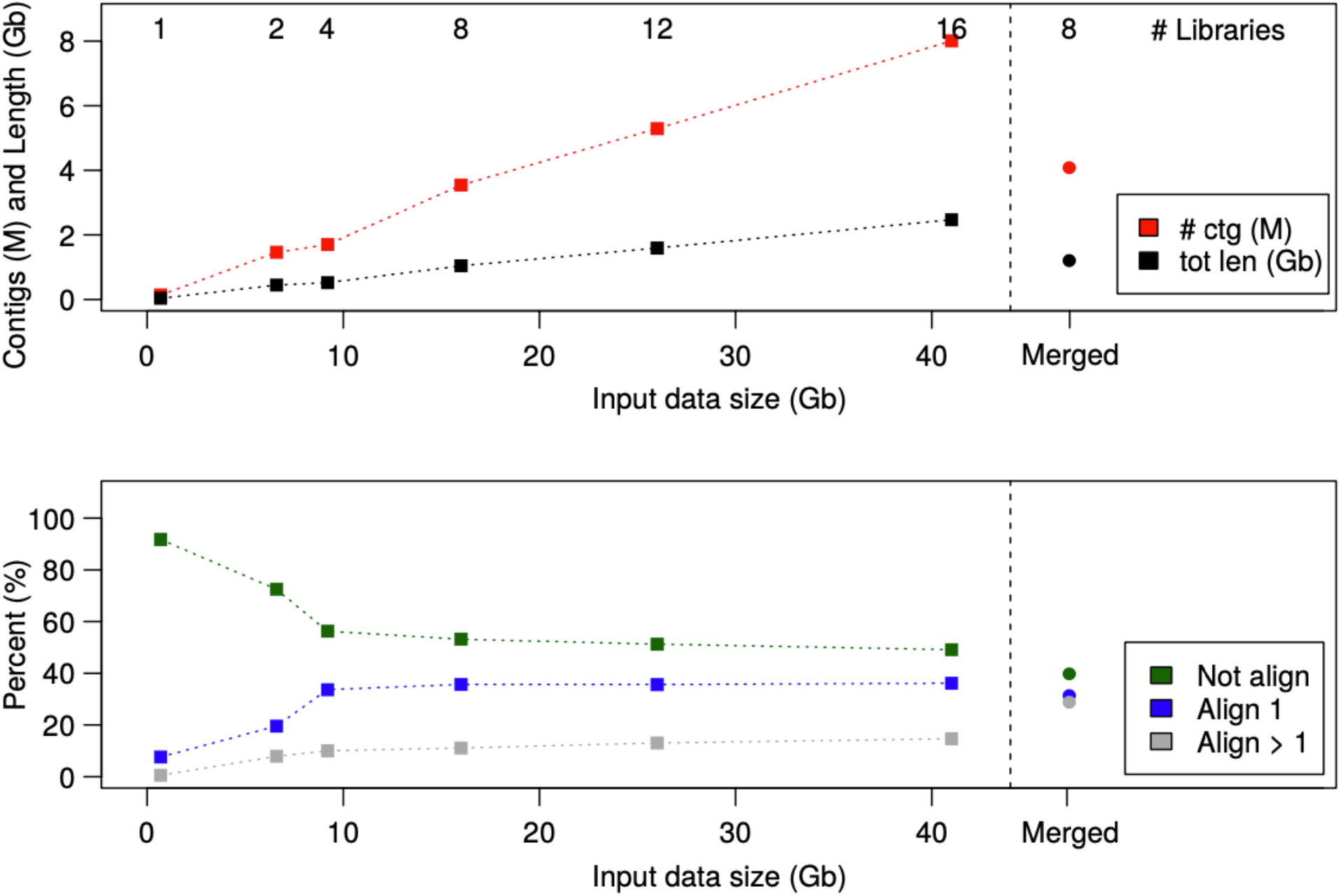
Assessment of reference metatranscriptome assemblies. (A) Total contig numbers (red) increase with input data size linearly, and so does total length of output assembly (black). The individual assembly of eight libraries that were subsequently merged together (shown to the right of the hatched line) has a similar number of contigs and length as the eight libraries assembled simultaneously. (B) The percentage of unaligned reads decreases as more libraries are added, initially with a rapid decrease until four libraries are added, and then with a more gentle slope as additional libraries are added. The benefit of increasing from eight to 16 libraries is not as evident in the percentage of reads aligned, suggesting that once four or eight libraries are assembled together, not much benefit is added by increasing the number of libraries in this dataset. Interestingly, the merged assembly has slightly fewer unaligned reads, but double the number of multi-mapping reads (align > 1, grey), indicating a strong amount of redundancy is still present in the merged assembly that is not present in the 16 library simultaneous assembly.

**Supplemental Data**

**Supplemental File S1.** Complete environmental and metadata for all samples.

**Supplemental File S2.** Overview of lineage data assigned to contigs.

**Supplemental File S3.** Full list of genera with significant linear models to pCO_2_ concentrations.

**Supplemental File S4.** Full list of genera with significant different abundances between early and late summer samples.

**Supplemental File S5.** Full differential gene expression analysis including genes differentially expressed between pCO_2_ levels and between seasons, as well as genes found in both comparisons.

**Supplemental File S6.** Full Gene Ontology analysis including GO enrichment for each differentially expressed gene list for Biological Process (BP), Cellular Component (CC), and Molecular Function (MF). Viral transcripts identified in BP in the season differential analysis are also included.

## Notes

https://doi.org/10.6084/m9.figshare.18128966.v1

### Competing Interest Statement

The authors have declared no competing interest.

### Summary of Updates

Reduced Introduction focus on oysters and increased focus on the effect of abiotic variables on microbial communities; included multiple test corrections for taxonomy by season or taxonomy by pCO2 level analyses; and revised (minor) for clarity.

